# CB-1 receptor agonist drastically changes oscillatory activity, defining active sleep

**DOI:** 10.1101/2024.07.18.604173

**Authors:** Irina Topchiy, Bernat Kocsis

## Abstract

Brain oscillations in different behavioral states are essential for cognition, and oscillopathies contribute to cognitive dysfunction in neuro-psychiatric diseases. Cannabis-1 receptor (CB1-R) activation was reported to suppress theta and fast gamma activities in rats during waking exploration, and here, we show that cannabis fundamentally alters network activity during sleep, as well. Prominent theta rhythm is present in rapid eye movement sleep (REMS), whereas fast oscillations appear as regular sequences of sleep spindles during intermediate sleep (IS) - both implicated in dreaming and memory consolidation. The CB1-R agonist disrupted these mechanisms, restructuring IS-REMS episodes; IS lengthened 6-fold and intruded REMS, where on-going theta was drastically reduced. The spindle architecture was also affected; its amplitude increased, and its peak frequency down-shifted into the theta range. Cannabis is known to induce psychotic-like conditions and cognitive deficits; thus, our results may help understanding the dual effect of cannabis on cognitive states and the role of network oscillations in psychiatric pathology.

## 1. Introduction

Network oscillations are essential for cognitive functions^1-3^, are modified by a wide variety of psychoactive substances^4-9^, and oscillopathies contribute to cognitive dysfunction in neurologic and psychiatric diseases^10-16^. Cortical spindles represent a unique member^17-20^ of the brain oscillatory hierarchy^21-26^; unlike other rhythms, they are short transients, appear during sleep and are generated by thalamocortical mechanisms rather than intrinsic cortical networks. Nevertheless, the cellular substrate of these oscillations and the mechanisms and the conditions of their functioning significantly overlap. Their role in cognition has been a primary target of intensive research for decades^27-31^, and spindle deficits have been firmly established as a robust endophenotypic marker of schizophrenia^32-34^.

Cortical spindles appear in slow wave sleep (SWS), become more frequent as approaching the SWS-to-REMS (rapid eye movement sleep) transition and are specifically enhanced in intermediate sleep (IS). IS is defined^17-20^ as short (<1min) episodes when delta rhythm, characteristic of SWS, vanishes in the electroencephalogram (EEG), and regular spindle sequences appear on the background of fully developed spectral characteristics of REMS, such as theta rhythm in the hippocampus (HPC) and low-amplitude high-frequency activity (gamma) in the frontal cortex (FrC). IS precedes every REMS episode, and it is the IS-REMS events that are primarily associated with sleep-related memory consolidation^35-40^.

Cannabis-1 receptor (CB1-R) activation has been reported to suppress theta and gamma activities in awake rats during exploratory behavior^8,9,^ but its effect on sleep oscillations has not been fully explored. To address this gap, we focused on IS-REMS episodes representing the active stages of sleep when the overall level of cortico-hippocampal activity reaches that in waking, in various cognitive tasks. We studied whether the psychoactive compound CP-55,940, a highly selective CB1-R agonist^41^, alters the structure of these episodes by acting on the unique sleep oscillatory pattern of thalamocortical spindles in the IS, which is immediately followed by HPC theta-dominated REMS. We observed a disruption of the IS-REMS; the IS lengthened 6-fold and appeared intruding into the REMS, which was entirely replaced by an “active sleep” segment with no standing HPC oscillations. The architecture of individual spindles was also affected. Unlike human EEG, where sleep spindles are readily recognized even by visual inspection, their detection in rodents based on spectral analysis remains difficult and rather ambiguous^42^. In this study, we employed a novel technique^43^, specifically adopted for spindle detection,^44-47^ which also allows examination of the fine composition of transient oscillations. Indeed, we found that sleep spindle architecture radically changes after CB-1R activation; spindles in the IS and those appearing in abnormal REMS are larger in amplitude, but their peak frequency becomes slower, shifting into the normal theta range. These findings suggest that meaningful spindle analysis has to go beyond counting spindles and include analysis of spindles’ inner structure and distribution relative to sleep stages not only to understand their role in cannabis action but also to resolve current uncertainties in connecting spindle deficits and cognitive dysfunction in psychiatric diseases, including schizophrenia^48,49^.

## 2. Results

Sleep-wake states, including the IS, were identified using standard criteria of FrC and HPC recordings and neck muscle EMG in 8 rats during 11-hour recording sessions before and after the injection of saline and the CB1 receptor agonist CP-55,940, with at least one week between the two injections (vehicle first in n=4 and CP-55,940 in the other n=4). Recordings were performed during the light period of the day (starting 8AM) when the rats were least active. The control vehicle (saline) was injected between the 3rd and 5th hours of the recording session, with the exact timing varying between experiments to separate the effect of the injection from potential circadian trends. The CB1 receptor agonist CP-55,940 was always administered in the third hour of the recording session, in a dose (0.3 mg/kg, intraperitoneally) which was shown effective altering rhythmic HPC activity during waking exploration^8,9^.

### Effect of CB1 receptor activation on IS-REM episodes

We found that the overall sleep-wake architecture was unaffected by CB1-R activation (Fig.1A), but the structure of IS-REMS episodes drastically changed (Fig.1B-D). IS segments, marked by spindle activity (11-15Hz), became longer whereas FrC gamma activity (30-50Hz) and HPC theta power (5-10Hz), i.e. the essential components of REMS, were severely suppressed. In control recordings (Fig.1C), in all rats (both before and after saline injection), wide-band delta (1-4Hz) activity dominated both FrC and HPC activities during SWS whereas during REMS low-amplitude fast gamma (30-50Hz) was present in the FrC and theta rhythm (5-10Hz) dominated HPC recordings. In the IS, REMS-like activity, i.e., HPC theta and FrC low-amplitude gamma signals, co-occurred with FrC spindles^17^, occupying an 11-15Hz frequency range (Fig. 1C). The length of REMS episodes stabilized by the second hour after the rats were moved to the recording cage in the morning and remained stable during the day (p=0.35; 147.1±14 and 144±11 s, respectively, in hr 2-3 and the last 2 hours of the control experiments). The average length of IS episodes was 26.0±1.9 s in the first 2 hours of recording (8-10AM) and slightly declined during the day (p=0.011; 12.9±1.5 in last two hours, 4-6PM), but saline injection did not have a significant effect on IS length (p=0.40, pre-vs. postinjection comparison) or on the number of IS-REMS episodes (p=0.15) (Suppl. Fig.S1).

**Figure 1.**
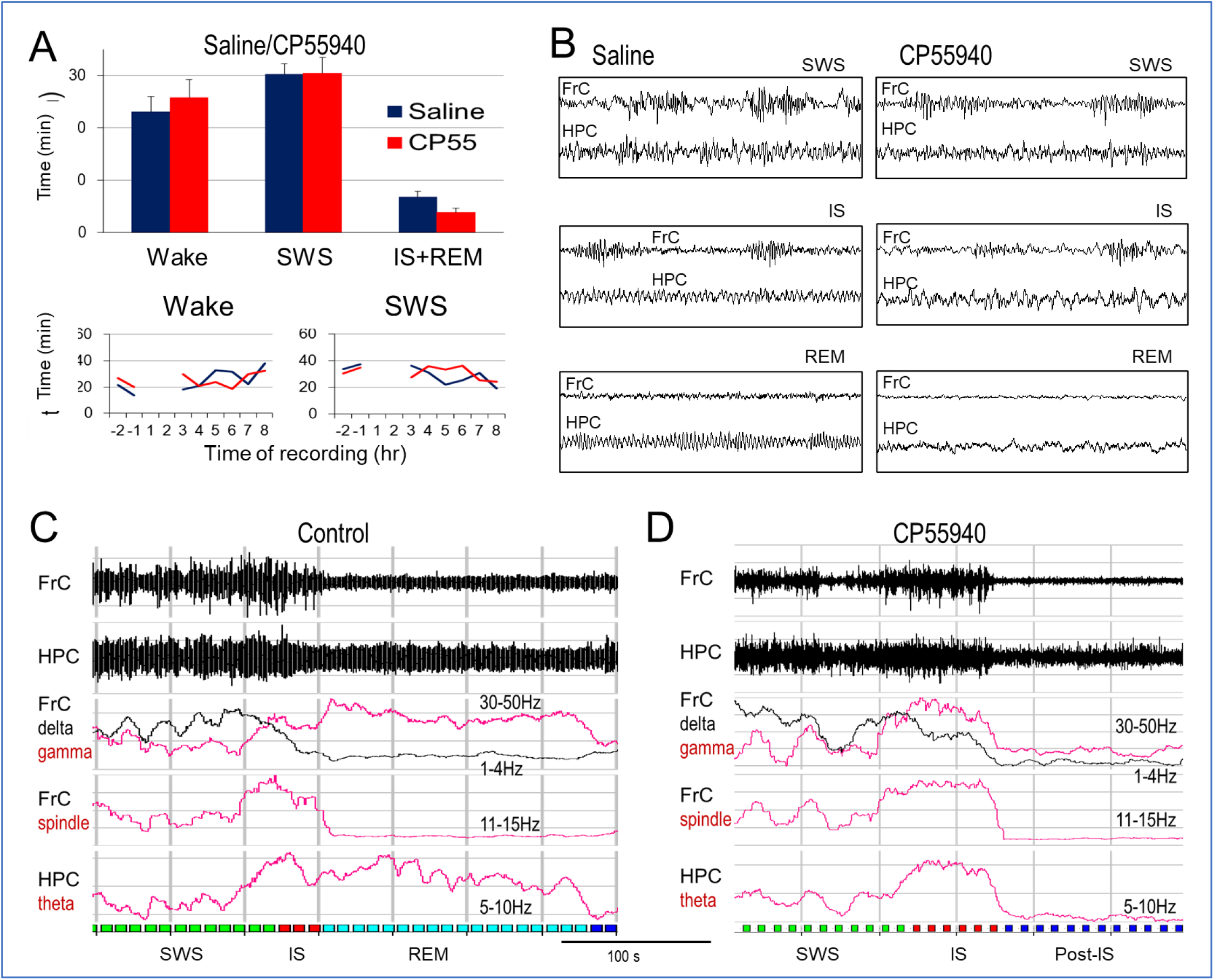
Effect of CB1 receptor activation on cortical and hippocampal activity in sleep. (**A)** Effect of CP-55,940 injection on sleep-wake architecture. T*op:* Average amount in sleep-wake states (AVE ± SEM) in the control group (saline) and after CP-55,940 injection. *Bottom:* distribution of sleep-wake states before (hrs -2,-1) and after (hrs 3-8) injection (*blue:* saline, *red:* CP55,940). Note that sleep-wake states disturbed by the injection procedure (hrs 1 and 2) are not shown. **(B)** Sample FrC and HPC recordings on an extended time scale (10s segments) after saline and CP-55,940 injection. **(C)** Sample FrC and HPC recordings through a SWS-IS-REM sleep episode in control and power distribution in different frequency bands (moving averages of filtered recordings) demonstrating drastic shifts in the oscillatory structure of FrC and HPC activity through these episodes. Note wide-band delta (1-4Hz) activity dominating both FrC and HPC activities during SWS switching to low-amplitude fast gamma (30-50Hz) in FrC and theta rhythm (5-10Hz) in HPC during REMS. In IS, REMS-like activity, i.e. HPC theta and FrC low-amplitude gamma, co-occurred with FrC spindles^17^, occupying an 11-15Hz frequency range. **(D)** Sample recordings and power distribution after administration of CP-55,940; note longer IS segments (*red* markers in the bottom trace) and reduced (apparent cessation of) FrC gamma and HPC theta power in post-IS period (missing *light-blue* markers).

No significant changes in the distribution of sleep-wake states were observed after CP-55,940 either; periods of “active” (including IS and REM) and passive (SWS) sleep alternated with wake states following the pattern normal for the light period of the day, similar to control recordings after saline injection (Fig. 1A). However, the structure of the IS-REMS episodes drastically changed after CB1 receptor activation. FrC gamma and HPC theta were drastically reduced in REMs (Fig. 1B), and the length of the IS substantially increased and appeared to substitute for REMS periods (Fig. 1D, 2B). The effect was robust, and long IS episodes were observed in all experiments (Suppl. Fig. S2). The average length of the IS episodes increased from 24.9±1.5s, starting in the first hour postinjection to reach a maximum of 111.7±20.6 s by the 8th hour (Fig.2A). The longest individual IS episodes (>80 s in all rats) were close to or even exceeded (>150 s in three rats) the average length of the control REMs (147.1±14; Fig. 1D). Due to the long IS, the rats spent more than three times longer in the state of IS than before the injection, even though the frequency of IS episodes remained stable after drug injection (Fig.2A).

Postinjection FrC autospectra were dominated by a sharp peak in the 5-10Hz range in all experiments, replacing the relatively wide band power (up to 15Hz) under control conditions (Fig. 2C). The dominant 5-10Hz component showed large interindividual differences; it drastically increased in 3 rats (p=0. 08 at the peak, 7-9Hz) and moderately increased in the others (p=0.22). However, the oscillatory character of the power spectra was still supported by the decrease in power in the adjacent frequency bands (1-5Hz and 10-15Hz, p=0.07 and 0.09, respectively) in all the experiments (see the autoscaled spectra, Fig. 2C).

**Figure 2.**
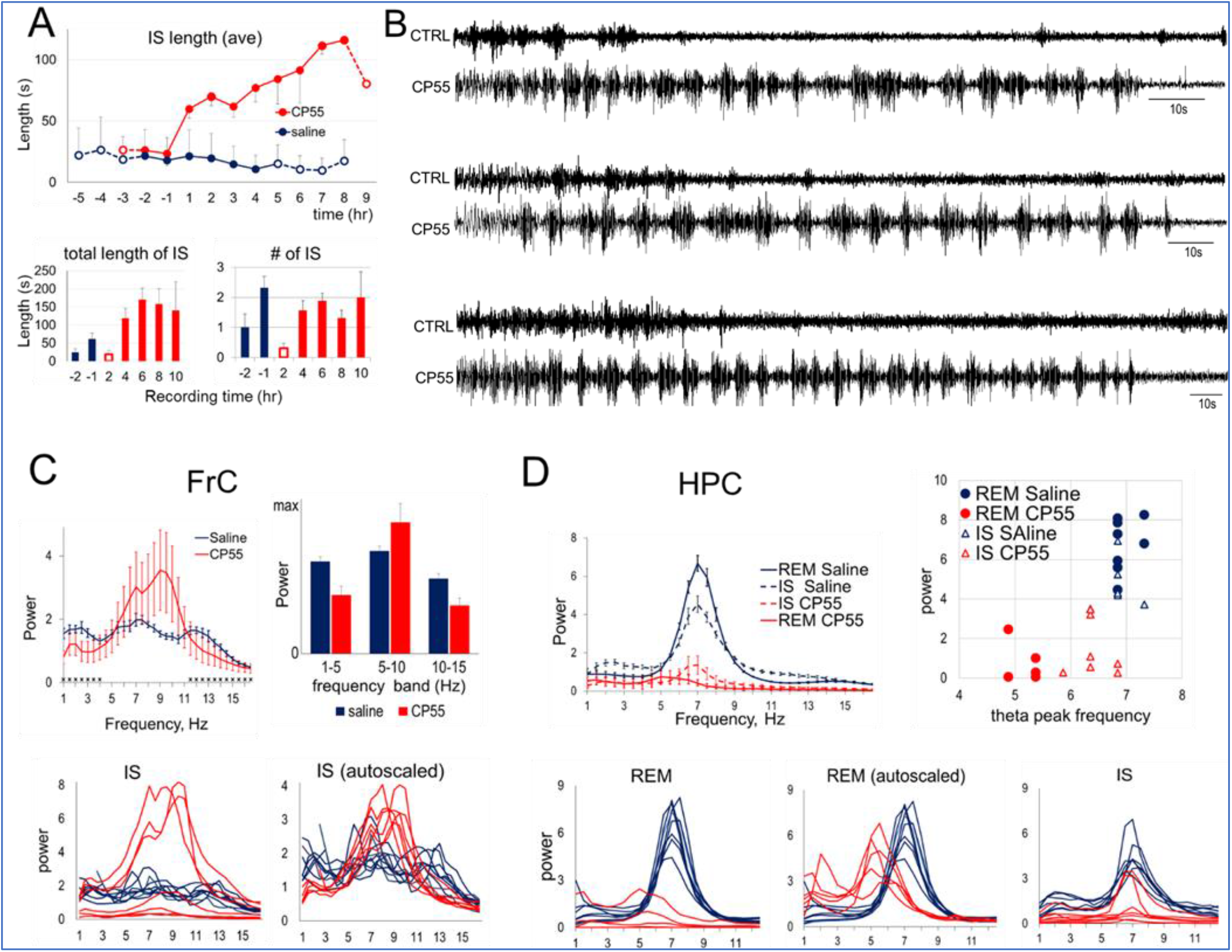
Effect of CB1 receptor activation on IS and REM sleep. **(A)** *Top*: Average length of IS episodes in 1-hr periods (injection at 0hr; n=8; open symbols: n<8, when either no IS was found in all rats or when no recordings were made in some shorter control experiments, see Fig.S1). *Bottom:* Total length and number of IS episodes before (hrs -2,-1; *blue*) and after injection (hrs 4-10; *red*). **(B)** Sample traces of FrC recordings during IS in three experiments before (CTRL) and after CP-55,940 injection (CP55). Note long IS episodes of varying lengths (time calibration: 10s), in all rats (see also Fig.S2). **(C)** Spectral analysis of FrC during IS. *Top*: Power spectra (mean±SEM; *left*) and power distribution in 5Hz frequency bands (*right*). Note dominant sharp peak in the 5-10Hz range, replacing wide-band power with three shallow “bumps” corresponding to delta, theta and spindle frequencies. *Bottom*: Autospectra in individual experiments, showing interindividual differences in peak power (*left*) and the autospectra autoscaled to reveal their altered character showing strong 5-10Hz peaks after CP-55,940 in all rats (*right*). **(D)** Spectral analysis of HPC during IS and REMS. Note drastic reduction of theta power in REMS as well as IS, and of theta frequency in REMS (6.96±0.07 to 5.17±0.11; p<0.001, n=5) but much less in IS (6.84±0.1 to 6.42±0.1; p=0.045), indicating potentially diverse mechanisms involved in generating the two rhythms. *Top*: Average HPC power spectra (mean±SEM) and theta peaks (power vs. frequency) in IS and REMS after injection of saline or CP-55,940. *Bottom*: Autospectra in individual experiments.

REMS episodes were difficult to detect, as one of its defining features, HPC theta activity markedly changed after CP-55,940 injection (Fig. 2D). Thus, in these experiments, we analyzed HPC power spectra in post-IS periods, which lasted 54.3±6.3 s, until they were disrupted by clear signs of waking (gross body movement in EMG) or SWS (high amplitude delta activity). No theta peak was present in the post-IS autospectra of HPC recordings in 3 rats, and it was significantly reduced in all others (p<0.001). In these experiments (n=5), the theta frequency also decreased, from 6.96±0.07 to 5.17±0.11 (p<0.001) (Fig. 2D). In IS, the amplitude of HPC theta oscillations was alsoreduced, and the frequency of oscillations decreased slightly after CP-55,940 injection (from 6.84±0.1 to 6.42±0.1; p=0.045). Yet, it was still significantly faster than postinjection REMS theta (p<0.001), indicating potentially diverse mechanisms involved in generating the two rhythms.

### Effect of CB1 receptor activation on spindle density and architecture

Closely following the temporal dynamics of IS length (Fig.2A), total spindle time increased after CP-55,940 injection due to a progressive increase in the number of spindles generated in each IS episode (p=0.001; 7.9±0.7 before and 21.3±1.8 after injection) and an increase in spindle length (p=0.023; 1.8±0.2s and 2.8±0.2s), whereas spindle density remained unchanged (p=0.076; 18.43±1.26 to 15.28±0.71) (Fig.3A). We next explored whether CP-55,940 had direct effect on the structure of the spindles, as well. Spectral analysis revealed a significant shift in the composition of IS episodes, dominated by a 5-10Hz component after injection (Fig. 2C). Low-frequency 6-9Hz spindles have been described in prior studies, but the limitation of FFT does not allow their clear separation from theta rhythm in the same frequency range.

Thus, spindle architecture was further analyzed using matching pursuit (MP), which allows decomposition of the signal recorded in the FrC during IS into a set of transient oscillations of different lengths, frequencies, amplitudes, and time locations (see details in Methods). It is an iterative procedure in which the signal is presented as a sum of Gabor functions (sinusoids enclosed in Gaussian envelopes), called “MP-atoms”, optimally chosen from a large and redundant dictionary^43,45^. We found that, besides their time-location corresponding to FFT-based spindle detection, MP analysis revealed consistent CP-55,940-induced alterations in spindle architecture, as well. These were represented by changes in the parameters of the MP-atoms, i.e., their center frequency and amplitude.

Figures 3B and 3C show examples of MP analysis of pre- and postinjection recordings. In the control recordings before injection, the centers of the MP atoms were in the 8-16Hz frequency range (Fig. 3B), confirming the basic spindle characteristics well documented in previous studies^17-19^. Postinjection, however, the centers of the MP atoms shifted to lower frequencies distributed in a narrower range below 10Hz (CP55 in Fig. 3B). The shape of the MP atoms also changed. In control recordings, spindles are represented by “sharp” vertically oriented components, corresponding to large within-spindle frequency variations. In fact, it is the rapid rise and fall of frequency within a short time that gives the characteristic shape of normal spindles. After injection, this dynamics was dampened, as depicted by shorter (i.e. truncated along the frequency (*y-axis*) MP-atoms, and even the appearance of “round” MP components, corresponding to increased spindle lengths (*x-axis*). MP analysis also detected longer, “horizontally oriented” (i.e. with homogenous intrinsic frequency) MP-atoms at lower frequencies, between 4-8Hz. The latter were present not only during spindles but also between spindles and during post-IS, most likely representing a standing (i.e., with no intrinsic frequency dynamics) HPC theta rhythm episodically transmitted to the FrC. In further statistical analysis, we applied filtering (see Methods) to eliminate these components (see *unfiltered* and *filtered* pre-injection recordings in Fig.3C). In control recordings, consistent with FFT-based spindle detection, MP-atoms appeared for only a short time (20-30 s) at the SWS-to-REMs transition, i.e., during IS (Fig. 4A, *top*). After CP-55,940 administration, however, spindles were present for at least 80s but increased to 150s during individual episodes (Fig. 2B). Importantly, although short episodes of transient theta acceleration occur in normal REMS (known as phasic as opposed to more stable tonic REMS), the slow spindles intruding in REMS after CP-55,940 injection maintain their characteristic pattern of rapid waxing-waning amplitude and frequency (compare Figs.3C, *top* and 4A, *bottom*).

**Figure 3.**
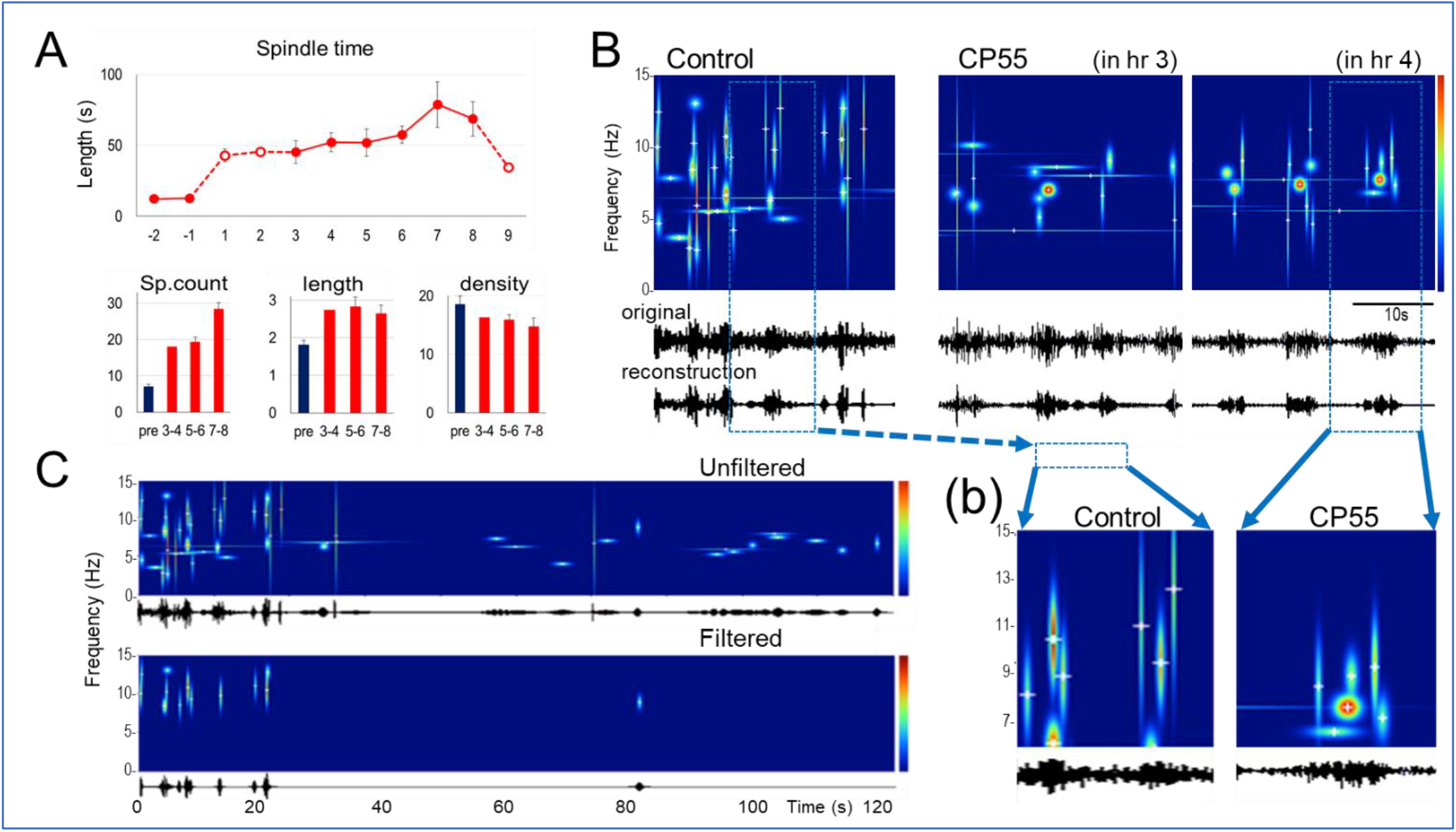
Effect of CB1 receptor activation on sleep spindles. **A**. *Top:* Total spindle time in 1-hr periods (N=8; open symbols: n<8 when no IS was found in all rats). *Bottom:* Number of spindles per IS episodes, average spindle length (in seconds) and spindle density (average number of spindles per minute) in IS episodes (2-hr averages) before (*blue*) and after injection (*red*). **B**. Effect of CB1 receptor activation on spindle architecture. Representative example of matching pursuit (MP) decomposition of FrC recordings into “MP-atoms” before (control) and after CP-55,940 injection (CP55; IS episodes) (25s segments, **(b):** zoomed 10s-sections). **C**. Filtering out long “MP-atoms” (>3s) in a preinjection FrC recording. Note that such filtering eliminates MP-atoms in the theta range during REMS (20-120s) with characteristics that are widely different from those of MP atoms in IS (0-20 s) and correspond to well-known characteristics of spindles. Theta rhythm in the FrC is normally transmitted from the HPC, and unlike on-going standing HPC theta, it appears with variable amplitude, which triggers the analysis procedure and generates MP atoms in the theta range (*unfiltered*). Thus, for statistical analysis, only “MP-atoms” with 0.5-3 s half-widths were selected within the 7-15Hz frequency band (*filtered*), whereas the amplitude threshold of spindle selection corresponded to the RMS criterion of spindle detection.

**Figure 4.**
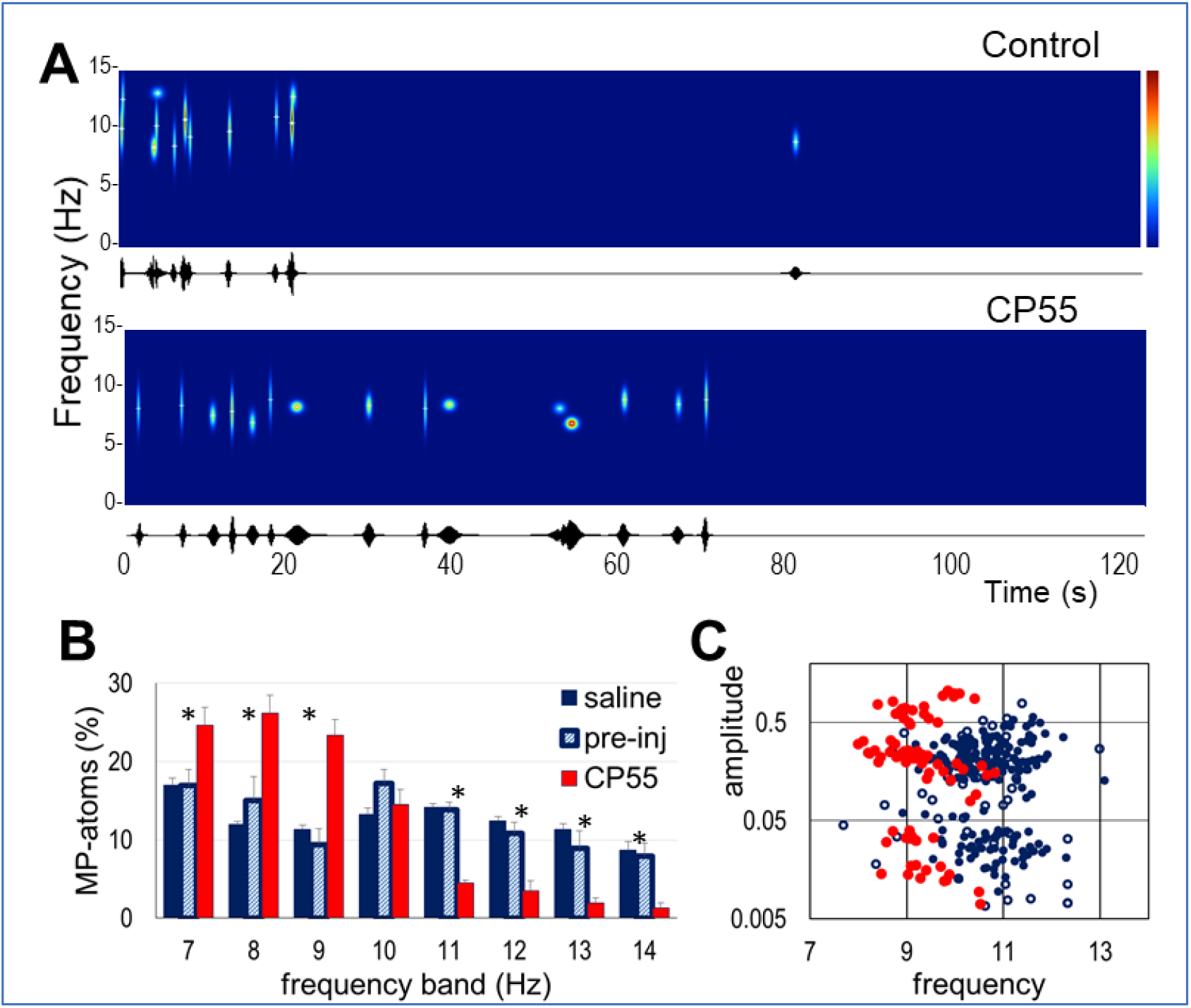
Shift in spindle frequency after CB1 receptor activation. **(A)** Example of MP analysis of 120s pre- and postinjection segments containing IS-REMS in control and IS—post-IS after drug injection. Note that ∼10Hz MP atoms are present only in the IS preinjection (0-20s) and that ∼7- 8Hz MP atoms extend over a long (70s) segment postinjection. The MP atoms were filtered to select the 7-15Hz and 0.5-3s half-width components to eliminate “horizontal”-oriented MP atoms representing segments short theta and only keep “vertical”-oriented MP-atoms representing spindles (see Fig.3C), shifted in the ∼7-8 Hz range. **(B)** Spindle frequency distribution in the control-saline and CP-55,940-test groups pre- and postinjection. *Y-axis*: distribution of spindles (MP-atoms) in 1Hz-frequency bands. * p≤ 0.05. **(C)** Amplitude-frequency distribution of spindles. Note the dominance of low-frequency (<10Hz) MP atoms postinjection (*red*) compared with faster (>10Hz) MP atoms preinjection (*blue, open circles*) and after saline injection (*blue, filled circles*) and the relative increase in the high-amplitude cluster after CP-55,940.

For group analysis of spindle architecture, IS bouts were subjected to MP decomposition, and average spindle frequency and amplitude were calculated for each IS bout, for each rat, and for each condition (a total of 331 IS bouts; 90 in the saline control, 49 pre- and 192 post-CP-55,940 injection). Thus, the MP atoms (0.5-3s long) of these IS bouts included both normal FrC spindles^18,19^ and altered spindles after CP-55,940 injection.

We found that CP-55,940 significantly changed the spindle architecture, particularly in terms of the dominant frequencies (<0.001). After injection, only 11% of all MP atoms had centers within the normal 11-15Hz range, while most (74%) had a center frequency below 10Hz (Fig. 4B, C). This finding was different from that of the control saline and preinjection recordings, where the distribution of MP-atom center frequencies showed two maxima, one corresponding to short theta segments during IS and the other (60% of all MP atoms) corresponding to normal spindles (*blue* in Fig. 4B, C). Spindle amplitude, the other key characteristic of MP-atoms, also significantly increased (p=0.015), showing two clusters of lower- and higher-amplitude spindles in both the saline control and CP5-55,940-injected groups (Fig. 4C).

## 3. Discussion

Our observations were remarkably consistent across experiments; selective activation of CB1-Rs fundamentally altered sleep spindle generation in all experiments, with no exception. Thus, the number of sleep spindles drastically increased after CB1-R activation, resulting in a 6-fold increase in the duration of IS, with intact spindle density and no change in general sleep-wake proportions. IS appeared intruding into REMS which was entirely replaced by an “active sleep” segment with no HPC theta (and/or FrC delta) oscillations. Individual spindle architecture also changed; spindle amplitude and length increased, and the peak frequency shifted from the normal range of 10-15Hz closer to theta frequencies (7-9Hz). Further investigations are necessary to clarify the underlying mechanism and implications of the solid findings presented in this report. This may be challenging. The robust increase in spindle activity in IS-REMS contrasts with the diminished power of continuous, i.e., nontransient, cortical oscillations at various (such as theta and gamma) frequencies induced by CB1-R activation (see also^50^). This finding is also in apparent contrast with reported deficits in sleep spindles in schizophrenia patients^32-34^ but more compatible with clinical situations in which exaggerated higher voltage spindles correlate with mental dysfunction^51^.

Cannabis is at the center of attention due mainly to two of its characteristic effects. First, it induces a transient, relatively short psychotic-like state, and second, it potentially has a lasting detrimental effect on cognitive function. The effects of CB1-R activation on IS-REMS shown in this study may be related to both factors. On the one hand, these sleep stages in humans are commonly associated with dreams—natural events with psychotic-like characteristics, labeled “normal delirium” by Hobson ^52^— and on the other hand, they may be involved in sleep-related memory consolidation^35,36^. The first has been traditionally associated with phasic events in REMS, such as eye movements, ponto-geniculo-occipital waves, and other transient patterns in the EEG. CB1-R agonist was shown to induce high voltage spindles in waking immobility and was suggested to be a potential mechanism of sensory “high” during recreational consumption of marijuana^50,^ although the translational value of the animal studies described here is obviously limited. In contrast, highly translatable cognitive tasks in rodents are well developed and are widely used to investigate the effects of CB1-Rs on different aspects of cognitive performance^9,50,53-57^. In addition to directly addressing CB1-R mechanisms at different levels (molecular, cellular, and in vivo pharmacological), the primary translational relevance of these investigations concerns psychiatric diseases in which cognitive dysfunction is present together with psychotic symptoms.

In human schizophrenia patients, impaired spindle activity^33^ is correlated with positive symptoms and deficits of sleep-dependent memory consolidation (rev.^34,58^). Thus, several studies have attempted to improve cognition in patients with schizophrenia by manipulating spindles. Recent studies have shown that eszopiclone significantly increases spindle density^59^; however, it fails to improve memory in both patients and healthy controls^60^. The negative outcome of this important and well-designed study is not necessarily inconsistent with the schizophrenia-spindle-memory association. This may indicate instead that the spindle count or spindle density alone may not be the most valuable biomarker representing causal relationships between schizophrenia and cognitive deficits. As demonstrated here using a psychoactive compound, the architecture of individual spindles is also subject to dramatic alterations—to the level at which specific parameters, such as peak frequency in our case, move this transient pattern out of its normal boundaries and into a range characteristic for a different class of oscillations (ongoing theta) while preserving other features, such as their waxing-waning burst character or density. Although existing analysis technology has limitations, some human studies have gone beyond calculating the number of spindles and reported deficits in spindle parameters, including amplitude, duration, density, and power^33,61-64^. This issue is not fully resolved, however. While some studies have reported decreases in both faster and slow spindles in schizophrenia patients^65^, others have shown deficits in fast-frequency spindles^66^ and reduced oscillatory frequency in frontal thalamocortical circuits^67^.

CB1-R agonists also have a marked effect on the system regulating the timing of sleep spindles. After CB1-R activation, spindles appear both in waking immobility^50^ and in REMS (our data), where they are normally absent. Reports of spindle deficits in human schizophrenia primarily concerned stage 2 (N2) non-REMS, and studies on the role of spindle activity in memory consolidation were also based on N2, with no reference to epochs immediately before REMS. However, abnormal spindle distribution relative to the REMS was recently reported in a large study of 11,630 individuals^68,^ which included an analysis of spindle density and frequency during “ascending” vs. “descending” N2, i.e., on the wake-N2-REMS vs. the REMS-N2-wake limbs of the sleep cycle, respectively. They reported a gradual (5 or more minutes) shift from slow (11Hz) to fast (15Hz) spindle activity preceding the transition from N2 to REMS but no change preceding the transition from N2 to wake (their Suppl. Fig. 16^68^). They also found a progressive (cycle-by-cycle) decrease in slow spindle density in descending N2 episodes and an increase in ascending N2 episodes over the night.

Direct analogy to rodent IS is not obvious in human EEG, where slow waves and spindles are abruptly suppressed within seconds of sleep state transitions^69^. However, signs of progressive non-REMS-to-REMS transitions are also present in humans. Polysomnographic (PSG) features of REMS appear gradually, starting several minutes before REMS onset with a reduction in muscle tone, followed by sawtooth waves, and terminating with bursts of eye movements^70^. The underlying neuronal mechanisms are under intensive investigation by combining PSG with stereoencephalography^71,72,^ providing further evidence that the non-REMS-to-REMS transition is a progressive phenomenon underpinned by local cortical sleep regulation. The state that might most likely be analogous to rodent IS is the 2-4 min period before REMS onset, starting in the lateral occipital and inferior temporal lobes, minutes before the frontal orbital cortex^71^. Local heterogeneity may also have a direct effect on the timing of global sleep state transitions. The HPC was shown to enter the REMS before the neocortex in humans^71,72^ as well as in rodents^73^. It was also shown that the delay between the scalp and intracranial EEG transition increased overnight, in parallel with the progressive dissipation of homeostatic non-REMS and increase in REMS pressure. Importantly, optogenetic inhibition of occipital cortex networks, i.e., the region to first exhibit REMS signs in humans^71^, suppressed the SWS-to-REMS transition in rats^74,^ indicating that diverse activity in local networks under homeostatic REMS pressure^75^ may be actively involved in switching global sleep states.

The mechanisms of neither dream generation nor memory consolidation are completely understood, but different aspects of the cellular network neurotransmitter mechanisms controlling sleep states and thalamocortical spindles have been studied in great detail, providing a large amount of data available for further investigation. We expect that extension of our study to cellular-molecular mechanisms may help in understanding not only the dual effect of cannabis on cognitive states but also the role of network oscillations, including sleep spindles, in psychiatric pathology.

## 4. Materials and Methods

Experiments were performed on male Sprague‒Dawley rats (250–300 g) under an animal use protocol approved by the Institutional Committee of BIDMC, in compliance with the Animal Welfare Act Regulations and with the Guide for the Care and Use of Laboratory Animals, National Institutes of Health guidelines.

### Surgery

Electrodes were implanted under ketamine/xylazine anesthesia (35–45 and 5 mg/kg of body weight, respectively) under aseptic conditions. Stainless steel jeweler screws were used to record cortical electroencephalograms (EEG) over the frontal cortex (FrC; 1 mm anterior and lateral from bregma) and for ground and reference electrodes over the cerebellum. Stainless steel Teflon-coated electrodes (125 µm) were implanted to record local field potentials in the hippocampus (HPC; AP 3.7, L 2.2, with one wire lowered 3.5 mm and another 2.5 mm from the surface of the skull). Two fluorocarbon-insulated 330-µm stainless steel wires were placed in the nuchal muscle on either side to record the electromyogram (EMG) signal. The wires were led to miniature connectors mounted on the skull by dental cement. To verify the position of the electrodes after the experiments, the rats were deeply anesthetized and perfused through the aortic arch with 0.9% NaCl followed by 4% paraformaldehyde (Sigma–Aldrich, Germany). The brains were removed, postfixed in 4% paraformaldehyde for 24 h and then stored in 30% sucrose. Forty-micron-thick sections were cut with a freezing microtome and stained with cresyl violet.

### Electrophysiological Recordings

Eight rats were used for chronic recordings starting one week after surgery. The recording was performed during the light period of the day (starting at 8AM) and lasted 11 hours. Either saline or CP-55,940 was injected IP at least 2 h after the recording started. The time of vehicle injection varied in hrs 3-5 to separate its effect from potential circadian trends (Fig. S1). The CB1-R agonist CP-55,940 was always administered in the third hour of recordings. CB1-R agonist CP-55,940 was administered at least one week before (n=4) or after (n=4) vehicle injection. Cortical and hippocampal EEG was recorded using monopolar electrodes referenced to the indifferent electrode placed over the cerebellum. Field potential recordings were filtered between 0.1 and 100Hz, with an additional analog notch filter at 60Hz, and sampled (with DasyLab 7.0, Microstar Laboratories, USA) at a rate of 250 samples/s simultaneously with EMG, which was filtered between 5 and 100Hz. The signals were then converted to Spike 2 (Cambridge Electronic Design, UK) for analysis.

### Materials

Animals were injected with the CB-1 agonist CP-55,940 (Pfizer, Groton, Connecticut), which was prepared as a suspension in methylcellulose (2.5 mg/mL). The injection volume of vehicle was 1 ml/kg. CP-55,940 was administered at a dose of 0.3 mg/kg.

### Sleep scoring

Sleep was scored in 10-s epochs, using standard criteria based on FrC and HPC recordings and EMG. Wakefulness was determined as a fast, low-amplitude EEG with concomitant high EMG tone. Slow wave sleep (SWS) was identified by increased slow irregular activity in EEG together with decreased EMG, and rapid eye movement sleep (REMS) was scored by the presence of standing, regular rhythmic activity in the HPC and an absence of EMG tone. Intermediate sleep (IS, transition from SWS to REMS) was determined when sleep spindles appeared in the cortex on the background of theta-activity in the HPC.

### Power spectrum analysis

Bouts of intermediate sleep (IS) and REM sleep of the entire recording, including pre- and postsaline or CP-55,940 injections, were submitted to analysis. Quantitative analysis of cortical and hippocampal EEG was performed by Fast Fourier Transform (FFT) using Spike-2 software. The power spectral density of the FrC EEG in the delta (1-4Hz) and gamma (30-50Hz) bands and the HPC EEG in the theta band (5-10Hz) were calculated and plotted over each recording. The power spectrum density of the EEG signals was calculated in 2.048 s segments (512 points, Hamming window) with a 0.49Hz frequency resolution. Absolute powers for separate IS and REM bouts were calculated in the range of 0.98 - 24.9Hz for each rat and were then divided to the averaged powers of this range for normalization. Additionally, power spectral data from CP-55,940 experiments were normalized relative to those of control (saline) recordings. The effect of CP-55,940 was assessed by a t test, p<0.05.

### Sleep spindle analysis

Cortical EEG was filtered between 7-15Hz and processed using Spike-2 software. The signal was rectified. Root mean square (RMS) was calculated with a time constant of 0.08. Subsequent smoothing of the signal was applied with a time constant of 0.08, using Spike-2 software. Sleep spindles during IS bouts were detected as RMS crossing the defined threshold, i.e. crossing the threshold by the RMS curve in ascending direction determined spindle start, and crossing by the descending part of the curve marked the spindle end. Individual thresholds were chosen for each rat and each condition (saline, pre- and postinjection of CP-55,940). Suprathreshold spindles 0.5 s and longer were selected for the analysis. Events with gaps of 0.1 s or less were amalgamated. Spindle quantity and length were estimated. The total time of spindling activity during IS was calculated and is expressed as a percentage for each condition. Spindle density was calculated as the number of spindles per minute expressed as a percentage. The effect of CP-55,940 was assessed by a t test, p<0.05.

Analysis of sleep spindle frequency was carried out with a matching pursuit (MP) algorithm using Svarog software (http://braintech.pl/svarog)^44-47^. MP is a procedure of an adaptive approximation of the signal to the functions chosen from a large set of waveforms “dictionary”. Waveforms in the “dictionary” are represented by Gabor “atoms” – sinusoids, modulated by Gaussian functions. The functions from this set that best represent the signal structure are chosen, and their weights are estimated. Thus, the MP presents the signal as a set of “atoms” in a time-frequency space. Each MP “atom” has a central point in time and frequency, and the time and frequency half-widths correspond to the mean ± σ on a Gaussian curve.

IS bouts in the saline control and pre- and post-CP-55,940 injection groups were filtered in the range of 7-15Hz, segmented into 10 s epochs, and decomposed to MP “atoms” with 100 iterations. A dictionary size of 99 “atoms” was used. The atoms with frequencies of 7-15Hz and 0.5-3 s half-widths were selected for analysis. The choice of the time and frequency range was determined both by analysis of frontal cortical spindles, which have slower frequencies^18,19^, and by an increase in spindle length and slowing spindle frequency at CP-55,940. The amplitude threshold for spindle selection corresponded to the RMS criterion for spindle detection. Averages of spindle frequency and amplitude were calculated for each IS bout, each rat and each condition (saline, pre- and postinjection of CP-55,940). The effect of CP-55,940 was assessed by a t test, p<0.05.

## Supplementary Figures

**Supplementary Figure S1.**
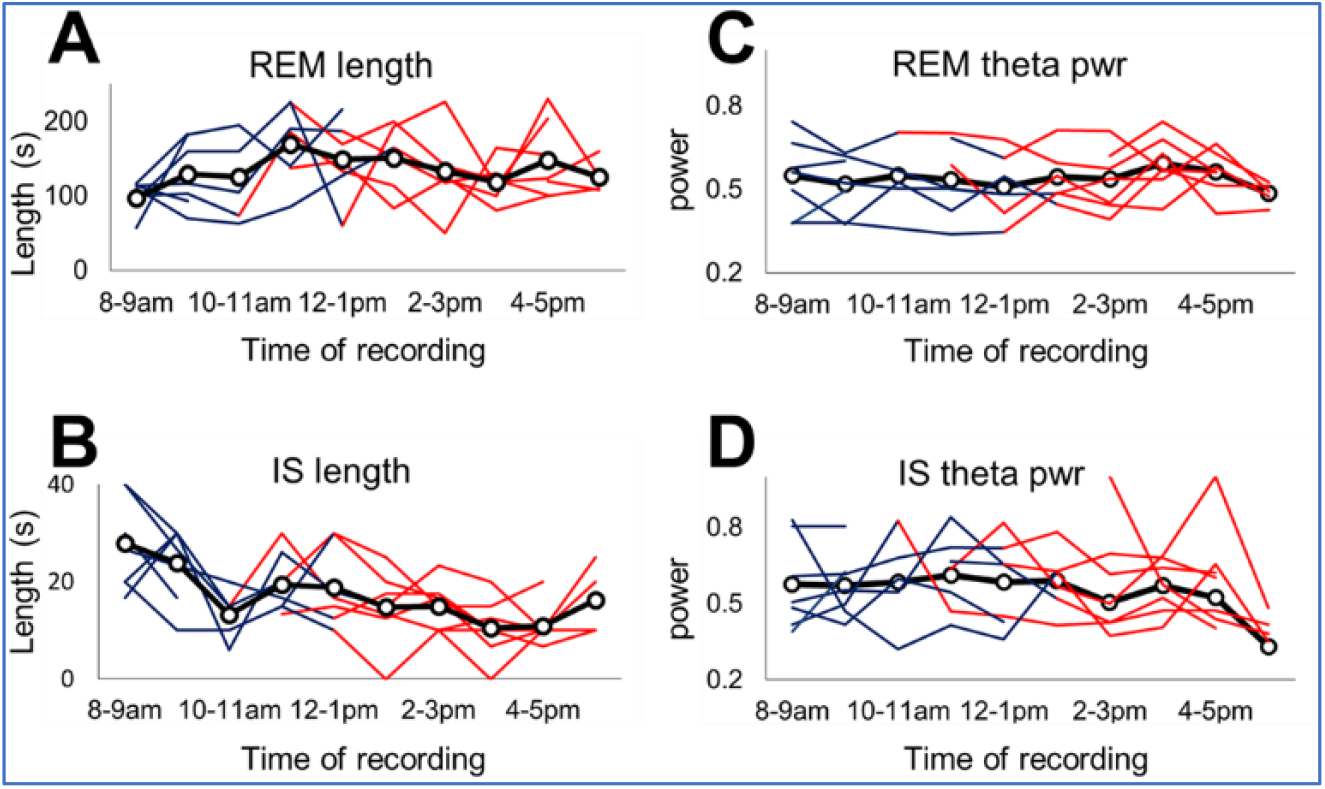
Hourly variation in the length of REM-S and IS episodes (**A-B**) and theta power in the HPC (**C-D**) in control experiments before (*blue segments*) and after (*red segments*) vehicle (saline) injection. (Group mean: **black—o—**). Saline was injected between the 3rd and 5th hours of the recording session, with the exact timing varying between experiments to separate the effect of the injection from potential circadian trends. HPC theta did not significantly change during the day (p=0.13 and p=0.36 for IS and REMS, respectively) or pre-postinjection (p=0.10 and p=0.23). Similarly, the 10-15 Hz power containing the spectral components of spindles (not shown) did not change either during the day (p=0.33) or in pre-postinjection comparisons (p=0.31).

**Supplementary Figure S2.**
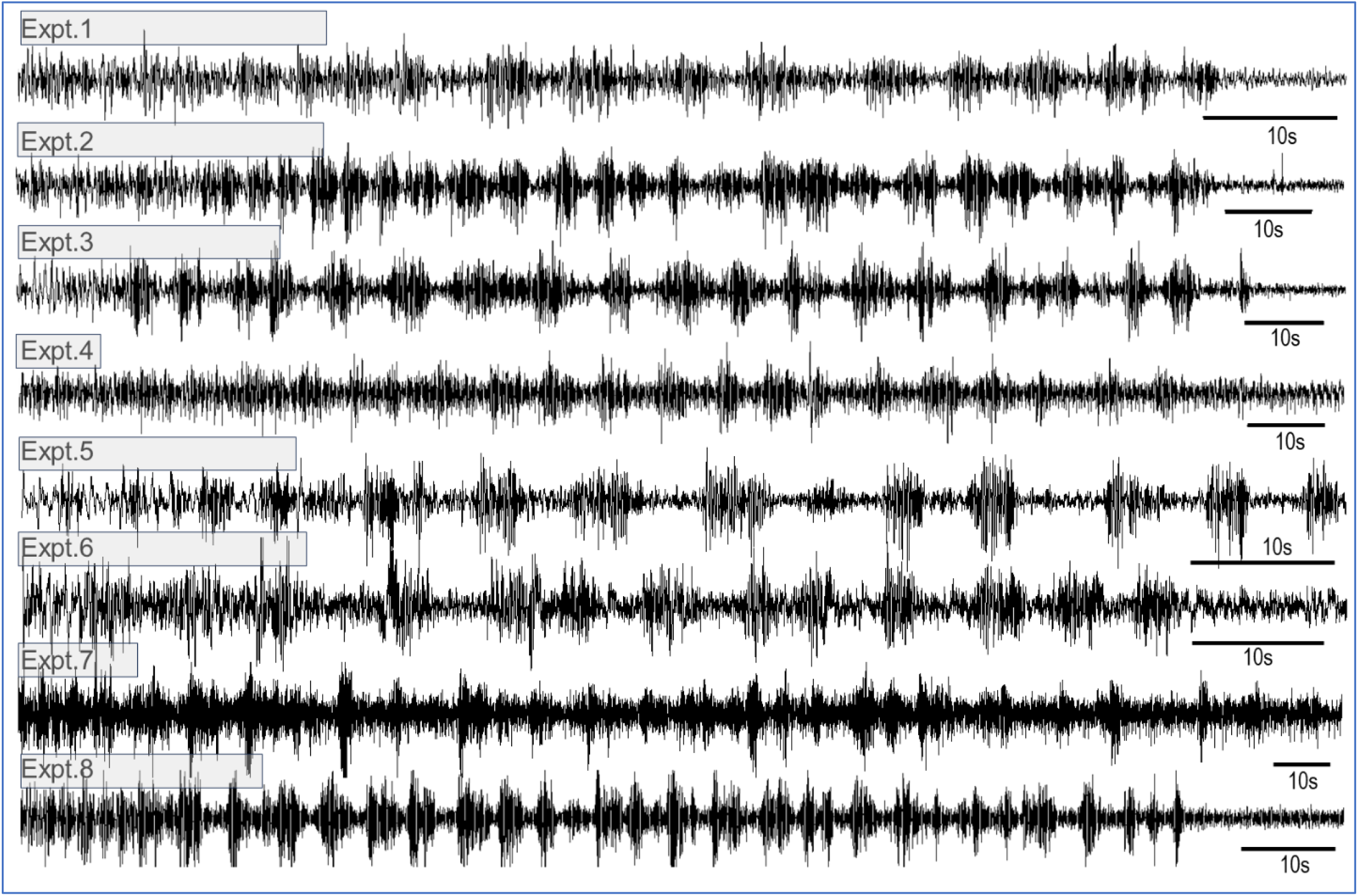
Sample recording of protracted IS episodes after CP-55,940 injection from different experiments. The effect was extremely robust; IS extension was a typical observation in all experiments (n=8), with no exception. The IS varied among the experiments (∼80s, 100s, 145s, 135s, 80s, 85s, 195s, 125s in these samples, in Expt.1..8, respectively; see time calibration: 10s), always exceeding the length of the control IS episodes several times (the frames around the titles indicating experiment numbers were calibrated to show the average IS length in control segments in each individual experiment).

